# Locust cGAS-like receptors recognize derivatives of a Gypsy retrotransposon to synergize with RNAi against viral invasion

**DOI:** 10.1101/2025.06.05.657997

**Authors:** Yao Xu, Yi-Lan Li, Ma-Cheng Zhang, Xin-Zheng Huang, Wang-Peng Shi, He-Ying Qian, Chuan Cao

## Abstract

The co-option of transposable elements (TEs) as immune sentinels represents an evolutionarily conserved strategy across metazoans, yet the molecular mechanisms linking retrotransposon reactivation to antiviral defense remain enigmatic. Here, we identify *LmGypsy*, a long terminal repeat (LTR) retrotransposon in *Locusta migratoria*, as a critical mediator of biphasic antiviral immunity against *Acrididae reovirus* (ARV). ARV infection triggers selective de-repression of *LmGypsy*, which orchestrates dual antiviral pathways: (1) its encoded reverse transcriptase synthesizes viral DNA (vDNA) from ARV RNA, fueling RNA interference (RNAi)-mediated viral RNA degradation through Dicer-2-dependent vsiRNA biogenesis; (2) LmGypsy-derived nucleic acids activate cGAS-like receptors (LmcGASs) and induce immune responses. Strikingly, vDNA persists in infected locusts from 24 h post-infection until host death, suggesting a role in sustaining antiviral activity akin to immune memory mechanisms observed in Diptera. This study provides the first evidence that insect *cGAS* homologs function as retrotransposon sensors, mirroring mammalian cGAS-STING recognition of endogenous retroviral elements. Our findings redefine retrotransposons as central hubs in arthropod antiviral immunity, bridging RNAi and nucleic acid-sensing pathways to establish a coordinated defense network. These results illuminate conserved principles of TE-immune crosstalk and highlight *LmGypsy* as a paradigm for understanding host-transposon coevolution in antiviral contexts.

## Introduction

Animal genomes, once conceptualized as static repositories of genetic information, are now recognized as dynamic systems shaped by evolutionary forces such as transposable elements (TEs)^1^. Transposable elements (TEs), once dismissed as genomic parasites or neutral “junk DNA,” are now recognized as pivotal drivers of genome evolution and functional innovation^2, 3, 4^. In vertebrates, TE domestication has generated novel regulatory networks and protein-coding genes, including components of placental development and immune pathways^2, 5, 6^. Similarly, insects exhibit TE-driven adaptations, including insecticide resistance in *Drosophila melanogaster* and climate adaptation through genomic plasticity^7, 8^. Notably, TE insertions disproportionately influence genes involved in stress response, behavior, and development, underscoring their role in phenotypic diversification^9, 10^. These observations highlight TEs as genomic “toolkits” that enable rapid adaptation to environmental pressures—a phenomenon termed *evolutionary tinkering*^11^.

The innate immune system, an ancient defense mechanism against pathogens, exhibits striking evolutionary interplay with TEs^12, 13, 14^. While host genomes have evolved stringent silencing mechanisms to suppress TE activity, accumulating evidence suggests that controlled TE reactivation during infection may serve as a strategic immune amplifier^13, 15, 16^. In mammals, retrotransposon-derived nucleic acids mimic viral PAMPs (pathogen-associated molecular patterns), activating sensors such as cGAS-STING to potentiate antiviral responses—a mechanism co-opted from TE-host arms races^15, 17, 18^. Drosophila studies reveal that retrotransposon-encoded reverse transcriptases convert viral RNA into immunostimulatory DNA, fueling RNA interference (RNAi) in insects^19, 20, 21^. However, the molecular logic underlying TE-mediated immunity—particularly in non-model insects—remains enigmatic, obscuring our understanding of how TEs balance their dual roles as genomic destabilizers and immune sentinels.

The migratory locust (*Locusta migratoria*), a devastating agricultural pest, offers a compelling model for investigating TE-immune interactions^22^. Its colossal genome (6.5 Gbp) harbors approximately 60% TEs, predominantly retrotransposons, making it one of the most TE-dense insect genomes known^23^. This genomic architecture, shaped by recurrent TE expansions, positions *L. migratoria* as a natural laboratory for studying TE dynamics and their functional consequences^24^. Our recent discovery of *LmGypsy*, a Gypsy-family retrotransposon selectively activated during reovirus infection, reveals a dual role for TEs in locust immunity. LmGypsy not only enhances RNAi efficiency by generating viral-derived small RNAs but also modulates the cGAS pathway to amplify antiviral signaling. These findings illuminate a sophisticated TE-host symbiosis, wherein ancient genomic “parasites” are co-opted as sentinels against viral threats.

This study provides the first evidence that insect cGAS homologs function as TE-specific sensors, mirroring mammalian cGAS-STING recognition of retrotransposons^15, 16^. By revealing how locusts exploit TE activation to establish a coordinated defense network, our work redefines retrotransposons as central hubs in arthropod immunity and highlights convergent evolutionary strategies across metazoans.

## Materials and Methods

### Locusts husbandry

Locusta migratoria eggs in this study were obtained from Hebei Province Locust Research Base (China). To eliminate potential ARV contamination, eggs were surface-sterilized with a formaldehyde-hydrochloric acid mixture (3% v/v formaldehyde, 1% HCl) for 5 min, followed by three rinses in sterile phosphate-buffered saline (PBS).

Eggs were incubated under aseptic conditions at 30 ± 1 °C and 70 ± 5% relative humidity (RH) until hatching. Newly emerged nymphs were maintained in controlled-environment chambers under standardized conditions (30 ± 1°C, 70 ± 5% RH, 16:8 h light:dark photoperiod) and provided with ad libitum access to freshly germinated wheat seedlings (*Triticum aestivum*). Developmental stages were synchronized by selecting newly molted 3rd-instar nymphs for all experiments

### ARV virions preparation

ARV purification was performed through ultracentrifugation with sucrose density gradient optimization. Briefly, Forty specimens of *Dasyhippus barbipes* (confirmed natural reservoir of ARV)^25^were homogenized in ice-cold phosphate-buffered saline (PBS, pH 7.4, Solarbio) using a sterile tissue grinder (Jingxin, Shanghai, China). The homogenate was centrifugation at 1,500 × g for 10 min at 4°C to remove tissue debris. Clarified homogenates were layered onto a pre-chilled discontinuous sucrose gradient (30%, 40%, 50%, 60% w/v in PBS) and subjected to centrifugation at 240,000 × *g* for 4 hr at 4°C in a SW 41 Ti rotor (Beckman Coulter, USA). Virus-containing bands at the sucrose interface were aspirated with spinal needles (BD Biosciences), diluted 1:10 (v/v) in sterile PBS, and concentrated through high-speed centrifugation (240,000 × *g*, 2 hr, 4°C). The resulting pellet was resuspended in PBS and aliquoted for storage at -80°C. ARV virions was verified through quantitative real-time PCR (qPCR) with the following pairs of primers: forward 5′-CTCCATCTCTCCGAAGTAACTC-3′; reverse 5 ′ -ACTGATCGATTGCGAGGTTC -3 ′ before proceeding with downstream applications..

### ARV infection and survival assays

Newly molted third-instar nymphs were intracoelomically injected with 9 × 10^12^ virions in 1 μL PBS using a Nanoject III microinjector (Drummond Scientific, USA). Control nymphs received equivalent volumes of sterile PBS (pH 7.4, Solarbio). Viral load dynamics were monitored by quantifying ARV genomic RNA in whole-body homogenates at 24 h intervals using qPCR with primers specific to the RNA-dependent RNA polymerase (RdRP) (forward: 5′-CTCCATCTCTCCGAAGTAACTC-3′; reverse: 5′-ACTGATCGATTGCGAGGTTC-3′). Survival assays were conducted over 27 days post-infection (dpi), with mortality recorded daily. All experiments included five biological replicates (n = 15 per replicate).

### RNA isolation and transcriptional profiling

Total RNA was extracted from whole nymphs using TRIzol reagent (Invitrogen) and treated with DNase I (Thermo Fisher) to eliminate genomic DNA. RNA integrity was verified via Bioanalyzer 2100 (Agilent Technologies). Strand-specific mRNA libraries were constructed from poly(A)-selected RNA (1 μg per sample) using the TruSeq Stranded Total RNA Kit (Illumina) and sequenced on a NovaSeq 6000 platform (2 × 150 bp; Novogene). Raw reads were quality-filtered using Trimmomatic v0.39^26^ and de novo assembled via Trinity v2.13.2.^27^. Differential gene expression analysis was performed using DESeq2 (FDR < 0.05, |log2FC| > 1)

### vDNA detection and small RNA sequencing

To assess retrotransposon-mediated reverse transcription of ARV RNA, total DNA was isolated from infected nymphs (n = 15 per timepoint) using the TIANamp Genomic DNA Kit (Tiangen). ARV-derived vDNA was quantified by qPCR using multiple primer pairs that span the entire ARV genome (Supplementary Table 1).

To determine if vDNA induces vsiRNAs, we inoculated 3rd-instar nymphs with 1 μg DNA from ARV-or PBS-infected locusts, and after 3 days total RNA from 15 nymphs was processed for next-generation sequencing of small RNAs. For small RNA profiling, 18–30 nt RNAs were enriched using the mirVana miRNA Isolation Kit (Thermo Fisher), sequenced on an Illumina HiSeq 2500 platform, and analyzed via the viRome pipeline after removing host-derived reads.

### Phylogenetic and functional analysis of cGAS-like receptors in *L. migratoria*

In mammals and *Drosophila*, cGAS is a cytosolic nucleic acid sensor that controls activation of innate immunity^28, 29^. To assess their role in locust antiviral immunity, putative cGAS homologs in *L. migratoria* (LmcGAS) were identified by BLASTp (e-value < 1 × 10^−5^) against reference sequences from *Homo sapiens* (Q8N884), *Sus scrofa* (I3LM39), *Mus musculus* (Q8BSY1), and *D. melanogaster* (A1ZA55, A8DYP7). Phylogenetic trees were reconstructed using IQ-TREE (1,000 ultrafast bootstraps) the maximum likelihood method^30^. In addition, the structurally related mammalian oligoadenylate synthetase 1 (OAS1) and male abnormal 21 (Mab21) were also aligned and constructed the phylogenetic tree. Functional domains were annotated via InterProScan.

### RNA interference (RNAi) assays

Double-stranded RNA (dsRNA) targeting *LmGypsy* reverse transcriptase and *LmcGAS* was synthesized in vitro using the T7 RiboMAX Express RNAi System (Promega). Primers containing T7 promoter sequences (Supplementary Table 1) were used to amplify template DNA from *L. migratoria* cDNA. Newly molted 3rd-instar nymphs were injected with 1 μg dsRNA (or dsGFP control) and incubated for 24 h prior to ARV challenge. Knockdown efficiency was confirmed daily using qPCR within 3 days after dsRNA inoculation.

### AZT Treatment

AZT (3′-azido-3′-deoxythymidine) is a nucleoside reverse transcriptase inhibitor used to suppress reverse transcriptase activity. To elucidate the role of retrotransposon-derived in locust antiviral immunity, we assessed the impact of AZT (Sigma-Aldrich) on the antiviral response of locusts following inhibition of reverse transcriptase activity. Newly molted third-instar nymphs were subjected to a 12-hour starvation period. Fresh wheat leaf segments (0.5 × 0.5 cm) were then coated with 20 μL of either 93 mM AZT or sterile PBS as a control. Subsequently, the nymphs were fed the AZT-or PBS-coated leaves for 24 hours before being inoculated with ARV virions. Viral load was quantified at 72 hours post-inoculation using qPCR.

### Statistical analyses

All statistical analyses in this study were performed using GraphPad Prism v9.5.^31^. Comparisons between groups for gene expression and viral loads were analyzed by one-way ANOVA and *t*-test. The standard error and sample size for each experiment are stated in the figure legends. Only *P*-value ≤0.05 were considered statistically significant. **Ethics Statement** This study utilized the migratory locust (*L. migratoria*), an invertebrate insect species. The experimental procedures employed (e.g., viral infection, behavioral observation, tissue dissection, etc.) are standard practices in insect pathology and virology research and are not considered to cause significant pain, distress, or harm beyond that inherent in standard laboratory rearing. Locusts used in this study were obtained from Hebei Province Locust Research Base (China).

## Results

### ARV-triggered activation of a Gypsy retrotransposon contributes to antiviral immunity in *L. migratoria*

Building upon our previous findings that ARV can establish persistent infections in *L. migratoria*^32^, we sought to comprehensively characterize the genes and immune pathways implicated in antiviral responses. Following infection of 3rd-instar nymphs with ARV, we performed transcriptome sequencing at 1 and 3 dpi. Remarkably, we observed a significant upregulation of a retrotransposon, identified as the Gypsy retrotransposon (*LmGypsy*), as early as 1 dpi (*P* = 0.0254; Fig. 1a). This finding was further corroborated by qPCR analyses, which revealed a dramatic increase in *LmGypsy* expression following ARV infection (*P* < 0.0001; Fig. 1b). This rapid induction of *LmGypsy* suggests its potential role in establishing antiviral defenses, aligning with emerging evidence that the reactivation of TEs in virus-infected hosts can contribute to antiviral immunity^33, 34^.

**Figure 1.**
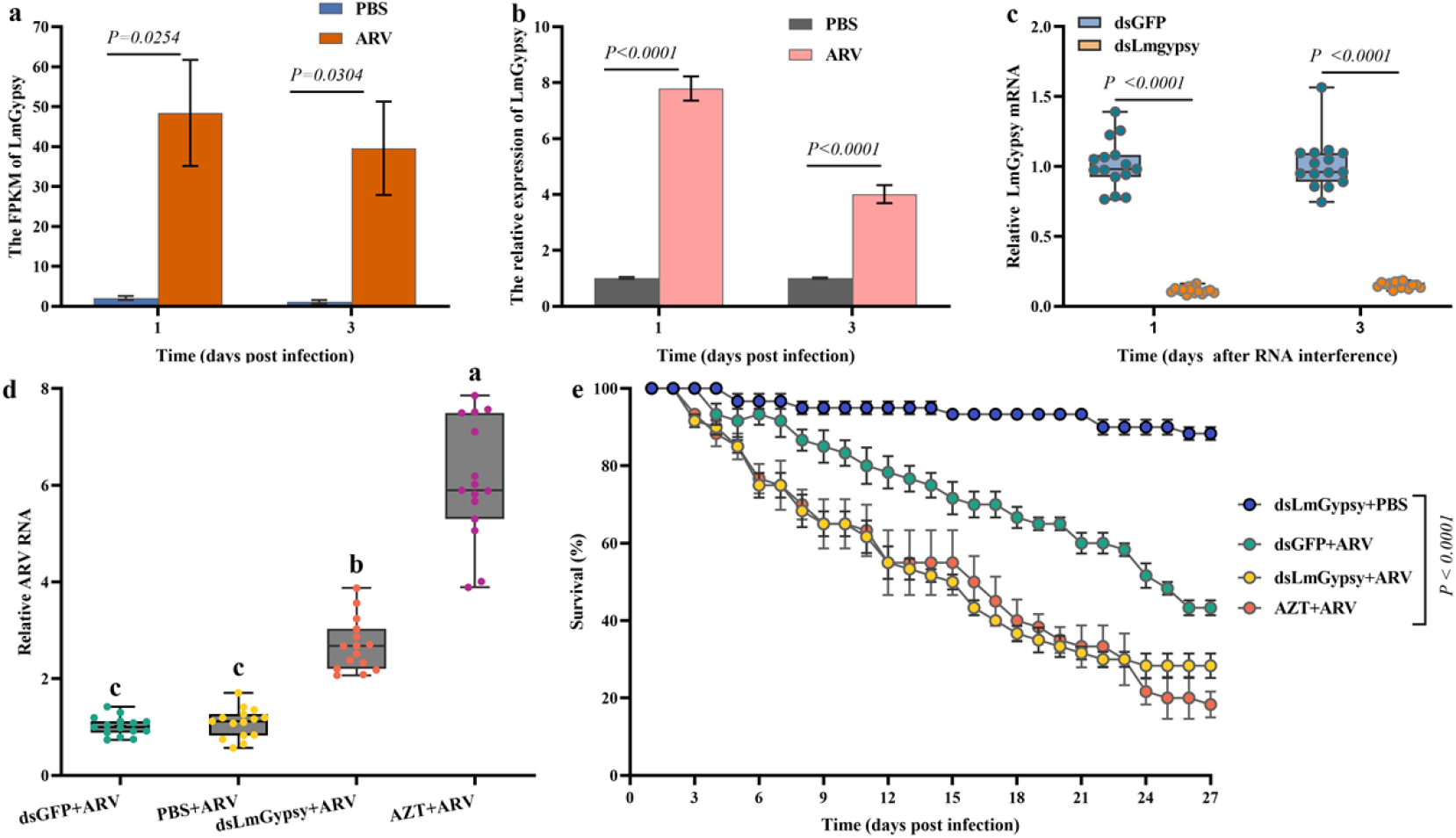
Activation of *LmGypsy* retrotransposon affects antiviral immunity and host survival in *L. migratoria* following ARV infection. **a**. Transcriptome analysis reveals significant upregulation of *LmGypsy* retrotransposon expression upon ARV infection. **b**. qPCR validation confirms a marked increase in *LmGypsy* expression following ARV infection. **c**. RNA interference targeting LmGypsy’s reverse transcriptase domain reduces its mRNA levels. **d**. LmGypsy knockdown or AZT treatment significantly increases the viral load in ARV-infected nymphs. **e**. Survival rates of ARV-infected larvae are significantly reduced following *LmGypsy* knockdown or AZT treatment. ARV infection assays were performed with five biological replicates, each contained 15 nymphs. The expression levels of *LmGypsy* or viral load were measured and indicated for every five nymphs.

To determine whether *LmGypsy* activation confers protection against ARV infection, we employed RNAi to specifically downregulate *LmGypsy* expression by targeting its reverse transcriptase domain. This targeted knockdown resulted in a significant reduction of *LmGypsy* mRNA levels (> 80%; *P* < 0.0001; Fig. 1c), while the knockdown nymphs developed normally without any observable phenotypic abnormalities. We subsequently assessed the impact of *LmGypsy* knockdown on viral replication. Nymphs infected with ARV and retaining *LmGypsy* expression demonstrated significantly lower viral titers at 3 dpi (*P* < 0.05; Fig. 1d). Conversely, nymphs subjected to *LmGypsy* knockdown were unable to effectively control viral replication, resulting in markedly higher viral loads at 3 dpi (*P* < 0.05; Fig. 1d). Furthermore, we evaluated the long-term effects of *LmGypsy* knockdown on host survival. Continuous rearing of nymphs for 27 days revealed that *LmGypsy* knockdown significantly reduced survival rates (*P* < 0.0001; Fig. 1d). Specifically, the survival rate in the ARV-infected control group was 43.33%, while the survival rate in the *LmGypsy* knockdown group decreased to 28.33%. (Fig. 1d). These findings collectively indicate that *LmGypsy* retrotransposon expression in ARV-infected locusts not only improves antiviral defenses but also significantly improves host survival.

In addition to these observations, we also assessed the effects of AZT treatment on ARV viral loads and locust survival. Treatment with AZT, a reverse transcriptase inhibitor, resulted in a significant increase in viral titers compared to untreated controls (*P* < 0.05; Fig. 1d), while simultaneously significantly reducing the survival rates of ARV-infected nymphs (Fig. 1e). This suggests the roles of *LmGypsy* and reverse transcriptase activity in modulating antiviral immunity and highlights its potential as a target for enhancing disease resistance in locusts.

### *L. migratoria* produces ARV-derived DNA via *LmGypsy* activity

In several species such as mammals, plants, and insects, non-retroviral RNA viruses can be reverse-transcribed into vDNA by retrotransposons or endogenous retroviruses^35, 36, 37^. However, it remains unclear whether a similar mechanism operates in other arthropods, particularly in Orthopteran insects such as *L. migratoria*. To investigate the presence of ARV-specific vDNA, we extracted genomic DNA from ARV-infected nymphs and amplified it using multiple primer pairs that span the entire ARV genome. All samples contained ARV-derived DNA, but only those originating from the complete RdRP coding sequence were identified, with this result further confirmed by Sanger sequencing. Using a pair of primers specifically designed for qPCR, we dynamically profiled vDNA synthesis kinetics, detecting vDNA as early as 24 h post-infection, with persistence observed throughout the infection period in locusts (Fig. 2a). This dynamic profiling underscores the rapid and sustained nature of vDNA synthesis in response to ARV infection.

**Figure 2.**
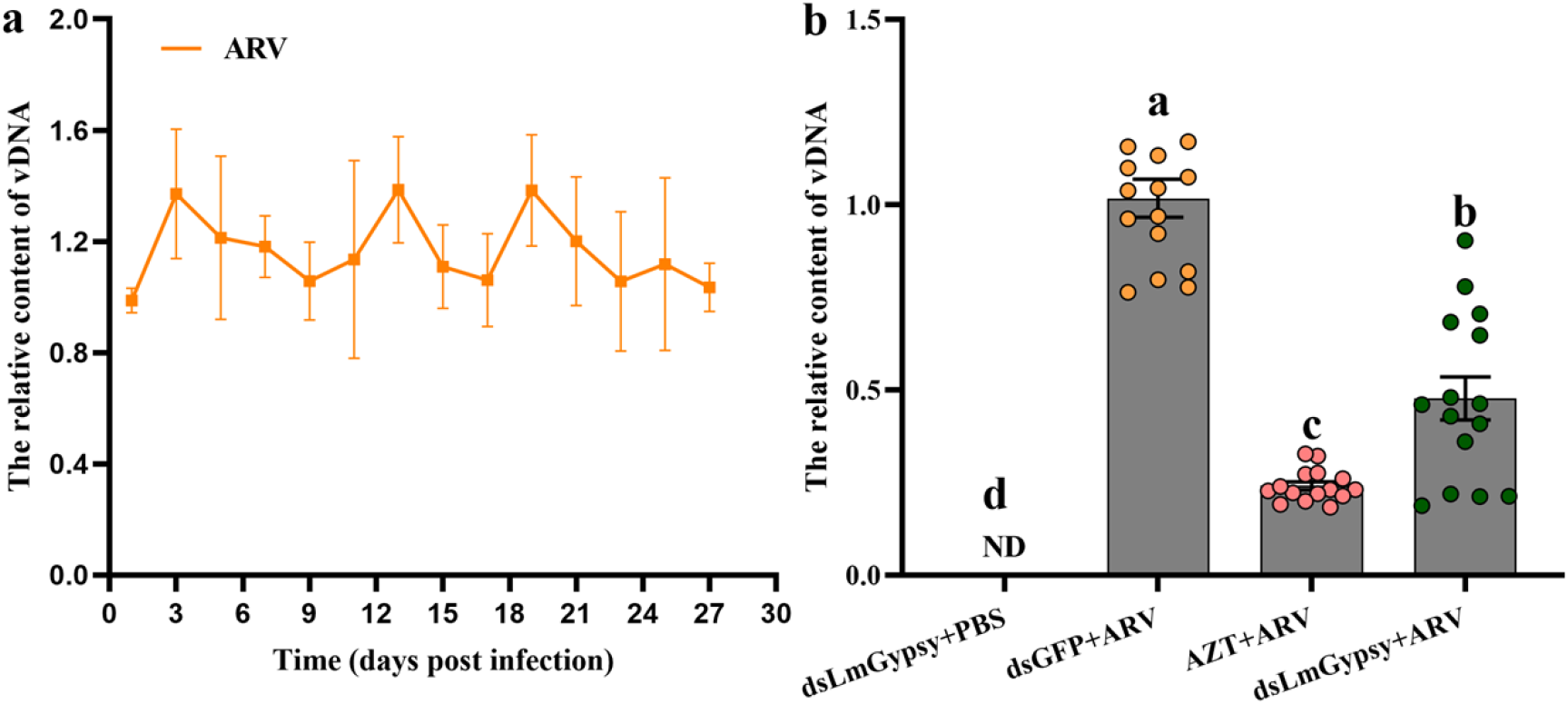
Biogenesis of vDNA during ARV infection in *L. migratoria*. **a**. Detection of vDNA present in ARV-infected locusts at various time points post-infection. no vDNA was detected in uninfected locusts, as indicated by the absence of data points for these samples. **b**. Impact of *LmGypsy* knockdown and AZT treatment on vDNA synthesis. Five biological replicates were performed. Each replicate contained 15 nymphs. ‘ND’ denotes that no vDNA was detected in these samples.

Given that retrotransposon-mediated reverse transcriptase activity has been implicated in vDNA formation in *Drosophila*, mosquitoes, and silkworms^19, 20, 38^, we hypothesized a similar mechanism in *L. migratoria*. Indeed, RNA interference-mediated knockdown of *LmGypsy* resulted in a significant reduction of vDNA production (*P* < 0.05; Fig. 2b). Furthermore, treatment with AZT further diminished vDNA synthesis (*P* < 0.05; Fig. 2b), reinforcing the role of reverse transcription in this process. Taken together, these results demonstrate that *L. migratoria* can generate ARV-derived DNA, and that *LmGypsy*-encoded reverse transcriptase is essential for vDNA synthesis.

### ARV-derived vDNA generates vsiRNAs that activate antiviral RNAi pathway

vDNAs have been reported to execute an antiviral function in insects^19, 20, 38^. To evaluate whether ARV-derived vDNA could protect locust from viral infection, third-instar nymphs were pre-inoculated with total DNA isolated from ARV-infected (vDNA) or uninfected locusts (LmDNA) 24 hours prior to challenge with ARV. At 72 hours post-infection with ARV, vDNA-treated nymphs exhibited significantly lower viral titer compared to LmDNA-treated controls (*P* < 0.0001; Fig. 3a). Therefore, our data provide compelling evidence for the role of vDNA in antiviral immunity of *L. migratoria*.

**Figure 3.**
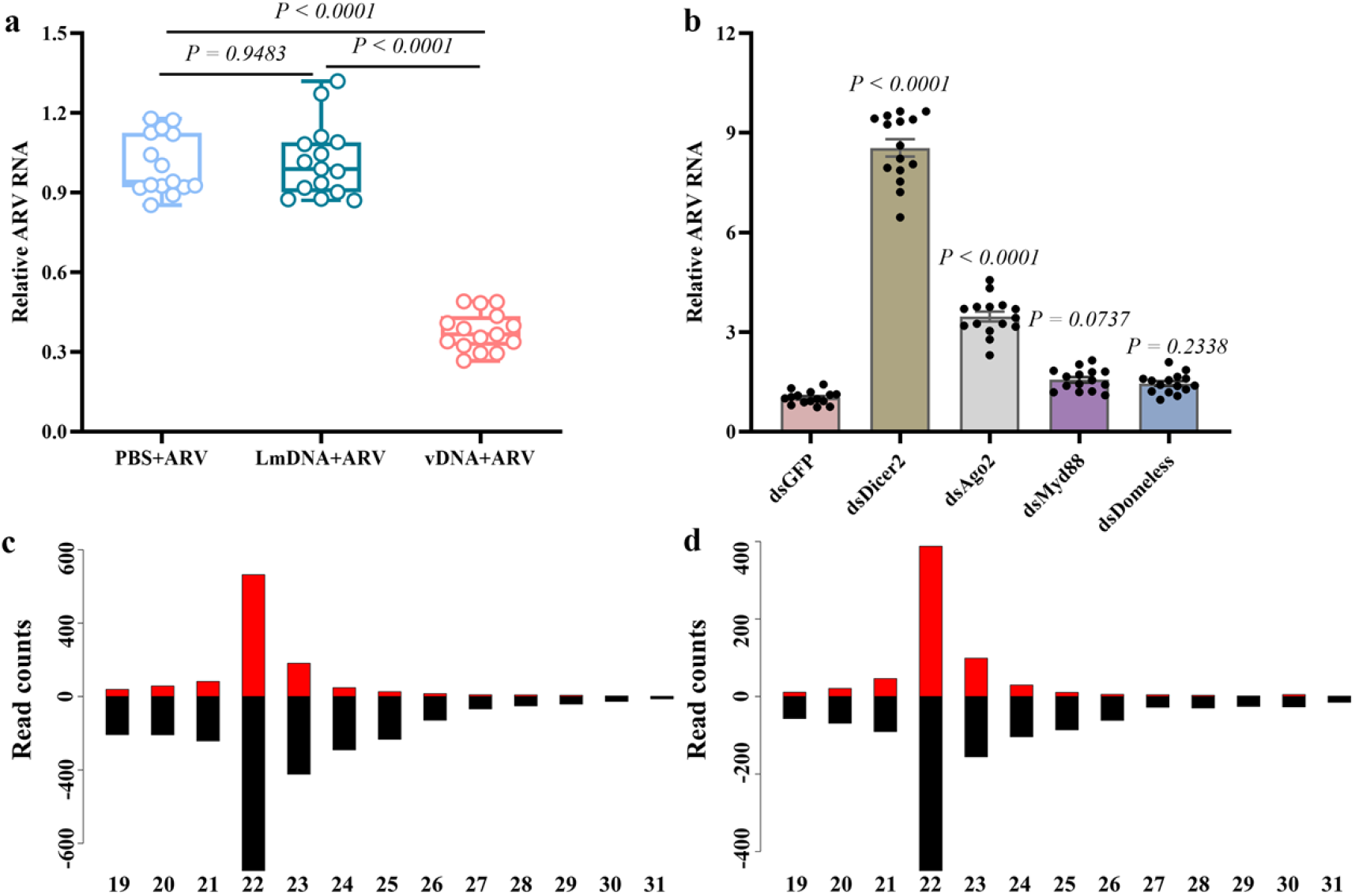
vDNA inhibits ARV replication through the RNAi pathway. **a**. ARV titers in vDNA-, LmDNA-, or PBS-inoculated locusts at 72 h post-infection. **b**. Effects of knocking down various immune pathway components on ARV replication. **c**. Size distribution of vsiRNAs from vDNA-inoculated locusts. **d**. Size distribution of vsiRNAs from ARV-infected locusts. ARV viral infection assays were performed with five biological replicates. Each replicate contained 15 nymphs.

Based on the findings of Poirier et al. (2018)^39^ and Zhu et al. (2022)^38^, it has been proposed that vDNA enhances the immune response of the RNAi pathway by boosting production of antiviral vsiRNAs. To define the mechanism underlying ARV-derived vDNA-mediated immunity in *L. migratoria*, we selectively knocked down key factors of immune pathways known for their antiviral roles in other insects^4^: *Dicer-2* and *Argonaute-2* (Ago-2) (RNAi pathway), *Myd88* (Toll pathway), and *Domeless* (JAK/STAT pathway). At 72 hours post-ARV infection, knockdown of *Dicer-2*, or *Ago2* led to a significant ARV accumulation in infected locusts (*P* < 0.0001; Fig. 3b). In contrast, *Myd88* or *Domeless* knockdown showed no significant effect (*P* = 0.0737; *P* = 0.2338; Fig. 3b), ruling out contributions from Toll or JAK/STAT pathways.

To investigate the contribution of ARV-derived vDNA to antiviral RNAi activation in *L. migratoria*, we delivered vDNA or LmDNA into nymphs and analyzed small RNA profiles 3 dpi. Small RNA sequencing revealed that vDNA-injected nymphs specifically generated ARV-derived siRNAs exhibiting characteristic features of antiviral RNA interference: (1) these siRNAs showed perfect strand complementarity with the RdRP of ARV; (2) they displayed a bimodal size distribution (19-31 nt) with a predominant peak at 22 nt, consistent with Dicer-2 processing of double-stranded RNA precursors(Fig. 3c)^41^. In contrast, LmDNA inoculation failed to induce virus-specific siRNA production. Notably, the length distribution and polarity pattern of vDNA-triggered siRNAs recapitulated the vsiRNA profile observed during active ARV infection (Fig. 3c,d), suggesting that viral DNA serves as a molecular signature to initiate Dicer-dependent antiviral RNAi responses in locusts. This finding provides direct evidence that vDNA sensing engages the canonical siRNA pathway to establish RNAi-mediated antiviral immunity in *L. migratoria*.

### LmGypsy expression can be sensed by cGAS sensors

Exogenous pathogens such as bacteria and viruses can induce host TEs expression^42, 43^. Once TEs expression reaches a critical threshold, cytosolic DNA sensors such as *cGAS* can detect these aberrantly expressed TEs and amplify immune signaling cascades^44, 45^. We hypothesized that *LmGypsy* expression might be sensed by cGAS-like receptors in locusts subsequently contributing to the immune response. To test this hypothesis, we conducted a search of the *L. migratoria* genome for homologs of known *cGAS* genes and identified four putative cGAS-like genes, which we designated *LmcGAS1* through *LmcGAS4* (Fig. 4a). Importantly, all four candidates encoded two conserved domains and contained a canonical active site compatible with cyclic dinucleotides (CDNs) synthase activity, suggesting functional conservation (Fig. 4a, 4b). Additionally, we also identified one Mab21-like protein (*LmMab21*) that was closely related to the cGAS-Mab21 subfamily (Fig. 4a, 4b). Phylogenetic analysis showed that *LmcGASs* cluster closely with *Drosophila* cGLRs (Fig. 4c), which function as pattern recognition receptors for sensing viral infections in flies^28^.

**Figure 4.**
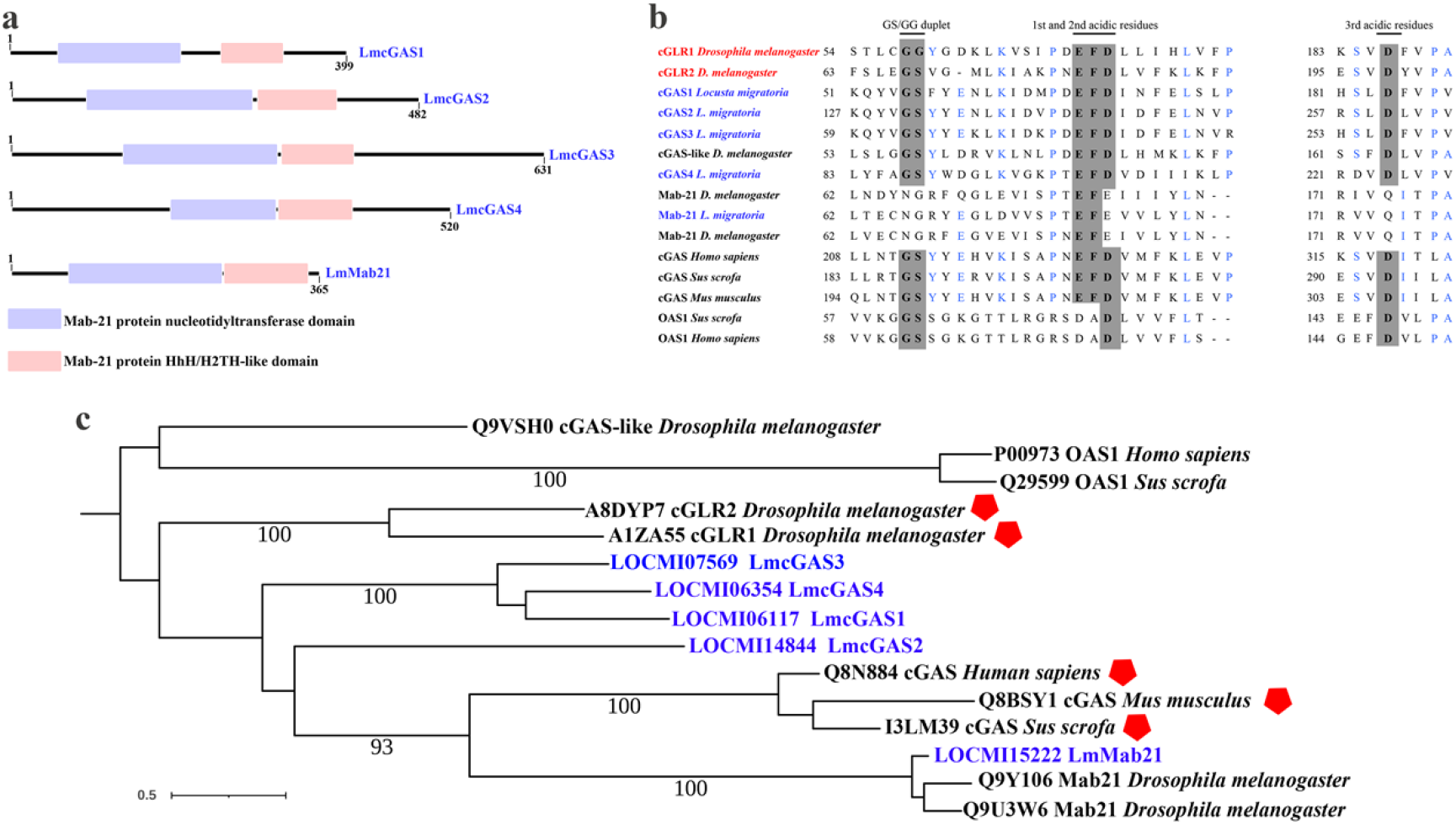
The cGAS candidates of *L. migratoria*. **a**. Domain organization of identified *LmcGASs* and *LmMab21*. **b**. Truncated alignment of identified *LmcGASs* and *LmMab21, Drosophila cGLRs* and *Mab21*, mammalian *cGASs* and *OAS1*. Essential active site residues (the GS/GG duplet and the metal ion coordinating acidic residues) were indicated above the alignment. **c**. Maximum likelihood phylogeny of the *cGASs, OAS1* and *Mab21*. cGAS proteins with experimentally validated roles in antiviral immunity are marked with red pentagons.

To delineate the functional roles of *LmcGASs* in antiviral defense, we individually knockdown the expression of *LmcGASs* and *LmMab21* by RNAi and measured ARV titers. Silencing *LmcGAS1-4* led to markedly enhanced ARV replication compared to the controls (*P* < 0.0001) (Fig. 5a), whereas *LmMab21* depletion showed no effect (*P* = 0.9990; Fig. 5a). Furthermore, ARV infection robustly upregulated *LmcGAS1-4* transcripts (*P* < 0.0001), but not LmMab21 (*P* = 0.7115; Fig. 5b), establishing these receptors as infection-responsive antiviral factors.

**Figure 5.**
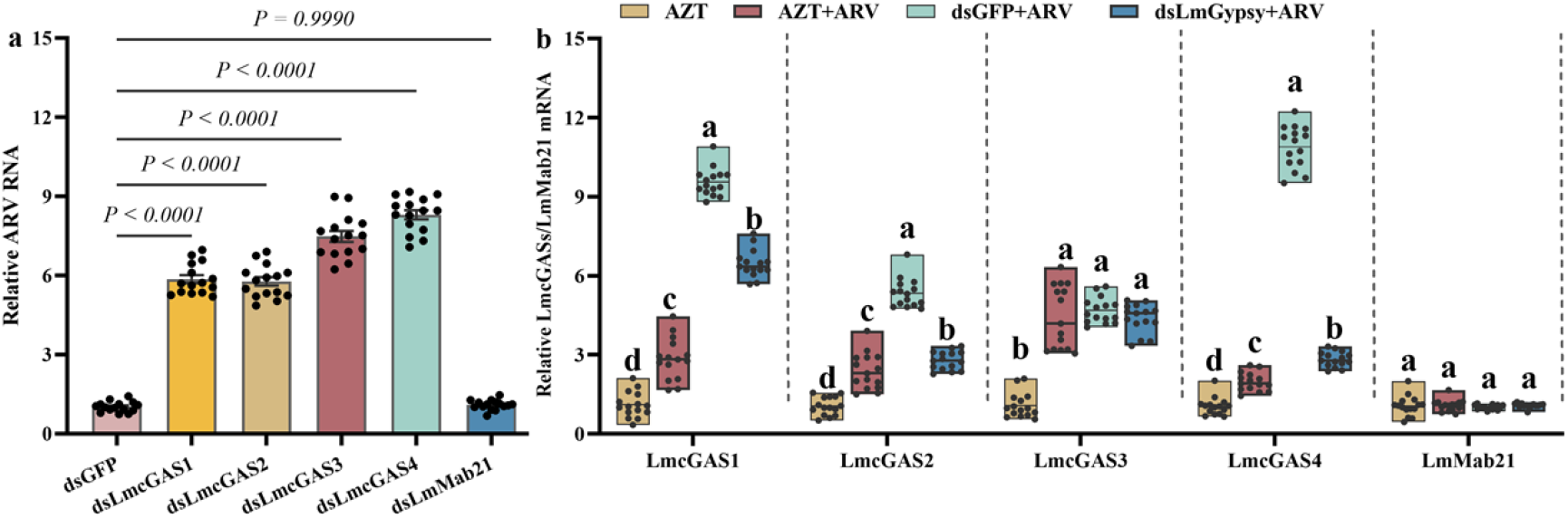
The antiviral response of *LmcGAS* genes is correlated with *LmGypsy* activation. **a**. Quantitative analysis of ARV replication following the knockdown of *LmcGAS1-4* and *LmMab21* expression. **b**. LmcGAS1-4 and *LmMab21* expression profiles in ARV-infected locusts subjected to *LmGypsy* knockdown or AZT treatment.

To investigate whether *LmcGAS* activation depends on TEs sensing—a mechanism conserved in mammals^45^—we analyzed *LmcGAS* induction under *LmGypsy* knockdown and AZT treatment. Strikingly, both *LmGypsy* suppression and AZT treatment abolished the induction of *LmcGAS1, LmcGAS2*, and *LmcGAS4* (*P* < 0.05), while *LmcGAS3* remained unaffected in ARV-infected locusts (*P* >0.05; Fig. 5b). These parallel outcomes demonstrate that retrotransposon-derived signals are critical for *LmcGAS1/2/4* activation during viral challenge.

## Discussion

### LmGypsy retrotransposon orchestrates a dual antiviral strategy through RNAi amplification and cGAS-like receptor activation

Our study establishes that *Locusta migratoria* employs a sophisticated two-pronged antiviral mechanism centered on the LTR retrotransposon *LmGypsy*, which bridges RNAi-mediated silencing and cGAS-like receptor signaling (Fig. 6). This work expands the emerging paradigm that TEs are not merely genomic parasites but are co-opted by hosts as immune amplifiers during infection^15, 42, 43^. Specifically, we demonstrate that ARV-induced *LmGypsy* activation generates both vDNA fueling RNAi responses and nucleic acid mimics that engage cGAS-like receptors (*LmcGASs*) (Fig. 6), revealing a functional convergence between insect and mammalian antiviral strategies.

**Figure 6.**
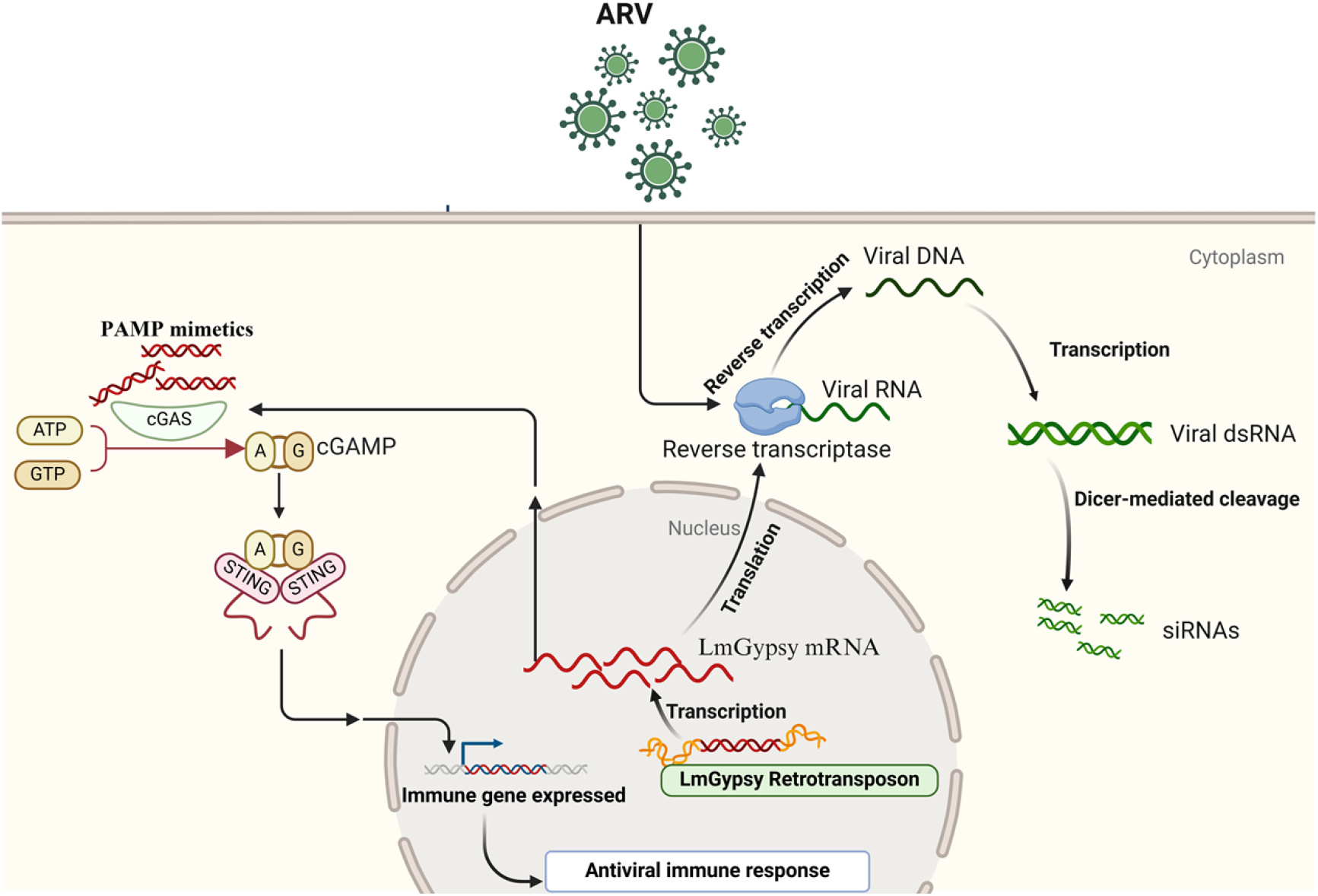
Proposed mechanisms of ARV-Triggered *LmGypsy* retrotransposon in antiviral immunity for *L. migratoria*. Two complementary mechanisms may explain how LmGypsy retrotransposons enhance antiviral responses in locusts: (1) *LmGypsy*-encoded reverse transcriptase converts ARV RNA to vDNA, which is transcribed to generate dsRNA, recognized and processed into vsiRNAs by the RNAi pathway. (2) Derepression of *LmGypsy* generates endogenous nucleic acids (e.g., dsRNA or retrotransposon-derived DNA) that mimic viral PAMPs. These “self-derived” signals are sensed by *LmcGAS* triggering downstream immune cascades.

### Evolutionary conservation of TE-mediated immunity

The discovery that *LmGypsy* retrotransposon activate *LmcGAS* receptors mirrors recent findings in mammals, where LINE-1 retrotransposons produce cytosolic DNA sensed by cGAS to restrict influenza A virus via the STING pathway^45^. Notably, our data suggest that insect cGAS-like receptors exhibit remarkable plasticity: while *Drosophila* cGLR1 senses dsRNA to produce 3 ′ 2 ′ -cGAMP^28, 46^, LmcGAS/2/4 in locusts respond to retrotransposon-derived ligands, potentially including vDNA or TE-specific dsRNA/DNA structures^47, 48^. This aligns with the broader hypothesis that cGAS-like proteins evolved as versatile sensors capable of detecting diverse nucleic acid patterns across taxa^49, 50^. The dependence of *LmcGAS* activation on *LmGypsy* (rather than direct viral genome sensing) underscores a layered defense strategy, wherein hosts exploit TE activity to amplify immune recognition—a mechanism likely under strong selective pressure given the prevalence of viral countermeasures targeting primary pathogen sensors^40^.

### vDNA as a nexus of RNAi and immune memory

The persistence of ARV-derived vDNA from 24 h post-infection until host death suggests a novel mechanism for sustained antiviral activity in insects. This phenomenon parallels observations in *Drosophila*, where RNA virus-derived circular DNA is vertically transmitted to progeny, conferring transgenerational immune resistance^51^. While our study did not assess heritable vDNA, the prolonged presence of vDNA in locusts raises intriguing questions about its role in immune memory. Furthermore, the detection of vDNA across diverse insects (e.g., *Bombyx mori* and mosquitoes) implies a conserved function in RNAi amplification^38, 39^, potentially compensating for the lack of interferon-based adaptive immunity in arthropods.

### TE derepression: a double-edged sword in host-pathogen conflict

ARV-induced LmGypsy activation aligns with reports that viral infections broadly dysregulate TE expression, as found in mammals infected with a variety of viruses, such as IAV, Zika, Ebola, and Rift Valley fever viruses, and in *Drosophila* challenged with *Drosophila C virus*, cricket paralysis virus, *Flock House virus* and invertebrate iridescent virus^14, 18^. However, the molecular triggers of TE derepression remain enigmatic. In mammals, SUMOylation loss of TRIM28 during influenza infection releases epigenetic silencing of endogenous retroviruses^52^, while in *Drosophila*, developmental cues license TE activation for future immune priming^14^. We propose that ARV may subvert host chromatin modifiers (e.g., histone acetyltransferases or DNA methylases) to activate *LmGypsy*, inadvertently triggering an “autoimmune” response against itself—a hypothesis supported by the exquisite sensitivity of *LmcGAS* receptors to TE-derived ligands. This paradoxical interplay underscores the evolutionary tension between TEs and hosts: while uncontrolled TE activity risks genomic instability, regulated activation provides a reservoir of immune-inducing signals^13^.

### Implications for insect-specific antiviral mechanisms

Our findings challenge the traditional view that insect antiviral immunity relies predominantly on RNAi. The convergence of LmGypsy-driven RNAi and cGAS pathways suggests a coordinated defense network. Notably, the functional divergence of *LmcGAS* isoforms (with *LmcGAS3* operating independently of *LmGypsy*) hints at receptor specialization, possibly for detecting distinct nucleic acid structures or subcellular localization cues—a feature observed in *Drosophila* cGLRs^28^. Additionally, the “PAMP mimetic” activity of *LmGypsy* products may explain how insects overcome viral evasion tactics, as TEs generate diverse ligands that are harder for pathogens to suppress compared to canonical viral PAMPs.

## Ethics and Compliance

All experiments complied with ethical guidelines for insect research. No human or vertebrate animal subjects were involved.

### Data availability

The raw reads of RNA-seq and sRNA data generated in this study are available at the NCBI SRA database with BioProject accessions PRJNA1206438 and PRJNA1206808. The nucleotide sequences of *LmGypsy, LmcGASs*, and *LmMab21* are available in Supplementary Table 2.

### Funding

This work was supported by the following grants:

National Key R&D Program of China (Grant No. 2022YFD1400500),

2024 Jiangsu Provincial Science and Technology Basic Research Program Youth Fund Project (Grant No. BK20241004),

China Agriculture Research System of MOF and MARA (Grant No. CARS-18-ZJ0101),

Research Start-Up Fund Project of Jiangsu University of Science and Technology.

